# Gravity-Dependent Metabolomic Responses in Starbor Kale *Brassica oleracea*: Comparisons With Simulated Microgravity Versus Gravity Grown

**DOI:** 10.1101/2025.09.29.679261

**Authors:** Sonia Kamran, Shagufta Perveen, Abubakar Salisu Barau, Yanru Li, Abdullahi Iro, George N. Ude, Supriyo Ray, Jie Yan, Anne A. Osano, Jiangnan Peng

**Affiliations:** Department of Natural Sciences, Bowie State University, Bowie, MD 20715, USA; Department of Chemistry, Morgan State University, MD 21252, USA

**Author notes:** Denotes equal contribution.

**Keywords:** Kale, Metabolomics, LC-MS, PCA, Microgravity, Gravity

## Abstract

As science continues to push the frontiers on the length of space flight, we begin to experience an enhanced need for food that is nutritionally dense. Nasa-grants have worked to explore the ability of growing superfoods in space on aircrafts. However, it is paramount to ensure the physiological health of the plants being grown in space flight, perform relatively similar to the plants that grow on earth. An aspect which needs to be further probed is the comparison of microgravity and gravity on the plants seeking to be grown in space flight. Within this NASA-funded research, we grew a superfood vegetable, Kale. The kale was grown in simulated microgravity environments to mimic life in outer space. This simulation was induced by means of a 2-D clinostat. Additionally, kale was grown in gravity conditions as well. We used multivariate statistical tools such as Principal Component Analysis (PCA), together with differential-expression visualizations like volcano plots, to summarize complex data rendered from LC-MS. In this study, we used these approaches to examine how horizontal and vertical orientations both in static and rotating configurations simulate aspects of simulated-microgravity and influence metabolic responses; this combined analysis provided a broad perspective on gravity-related metabolic adaptations and points to potential molecular markers that could guide future research on spaceflight health and countermeasure development.

## Introduction

As science continues to push the frontiers on the length of space flight, we begin to experience an enhanced need for food that is nutritionally dense. Literature underscores the importance of nutrient-rich diets for this in space flight, and the consequences of inadequate diets (Zhao, 2024; Baba et al., 2020; Morales-Navas et al., 2023; Smith et al., 2021; Douglas et al., 2022; Bourdier et al., 2022). While current food sources consumed during space flight are invaluable; there is a desire to provide astronauts with food equivalent to the nutrient content and bioavailability as found in standard superfoods grown on earth. With the effects of radiation being encountered by those in space flight, we turn to metabolites in plants to combat the effects (Johnson, 2003; Poulose et al., 2014). Flavonoids, a subclass of polyphenols, which are commonly found in superfood vegetables, are deemed to be effective in counteracting the effects of radiation which are experienced while in space (Nagpal, 2017; Dulcich, 2013). Nasa-grants have worked to explore the ability of growing superfoods in space on aircrafts. However, it is paramount to ensure the physiological health of the plants being grown in space flight, perform relatively similar to the plants that grow on earth.

An aspect which needs to be further probed is the comparison of microgravity and gravity on the plants seeking to be grown in space flight. Over the past five years a great deal of research has been accomplished in this area (Villacampa et al., 2021; Oluwafemi, 2021; Kruse et al., 2020; Medina et al., 2021), but further studies are needed to form a comprehensive and conclusive picture regarding the impact of microgravity on the health of growing plants. Spacial limitations have been placed on utilization of the international space station (ISS) for such experiments. Research has indicated that apparatuses such as clinostats can be useful in simulating microgravity conditions (Hoson et al., 1997; Wang et al., 2016). Within this NASA-funded research, we grew a superfood vegetable, Kale. The kale was grown in simulated microgravity environments to mimic life in outer space. This simulation was induced by means of a 2-D clinostat. Additionally, kale was grown in gravity conditions as well. We used multivariate statistical tools such as Principal Component Analysis (PCA), together with differential-expression visualizations like volcano plots, to summarize complex data rendered from LC-MS. In this study, we used these approaches to examine how horizontal and vertical orientations both in static and rotating configurations simulate aspects of simulated-microgravity and influence metabolic responses; this combined analysis provided a broad perspective on gravity-related metabolic adaptations and points to potential molecular markers that could guide future research on spaceflight health and countermeasure development.

## Materials and Methods

### Materials and reagents

LC-MS grade of methanol, ethanol, and formic acid were purchased from Fischer Scientifics.

### Cultivation of kale under microgravitational conditions

Starbor variety kale seeds had been purchased from West Coast Seeds Inc. The kale seeds had experienced germination in rockwool support. The seeds were watered for 3 weeks’ time.

Subsequently, 12 seedlings were transferred to modified Bio-World GA-7 “Magenta” growth vessels. These vessels were filled with soil for growth of kale. We simulated microgravity for the experimental kale by means of the 2-D clinostats set to 10 rpm, which we purchased through COSE (insert citation from paper A). Both the experimental and control plants were grown in a Conviron ATC26 growth chamber for 6 weeks. The temperature of the growth chamber was kept at 22.5°C, with CO_2_ levels reported to be between .06-.08%. Freshly collected kale samples were immediately frozen and lyophilized. The dry samples were stored in a freezer until LC-MS/MS analysis.

### Sample extraction

The lyophilized kale samples were ground. Three solvents were tested for the extract efficiency, including methanol, absolute ethanol, and 70% methanol in water. The kale powder (100 mg) was added to 5.00 mL of the solvent and sonicated for 30 min at 30 ^0^C. The extract was kept at room temperature for 30 minutes and then centrifuged at 3000 rpm for 5 minutes. Filter the supernatant through a 0.22 µm syringe filter. Transfer 100 µL of the filtrate to an HPLC vial and add 900 µL of methanol for the LC-MS/MS analysis.

### LC-MS/MS analysis

An Agilent 6546 LC-Q-TOF system equipped with an DAD detector and an autosampler was used in this analysis. An Agilent Eclipse C18 column (4.6 × 150 mm, 5.0 µm) was used for the separation. The gradient elution of 0.1% formic acid in water (solvent A) and 0.1% formic acid in acetonitrile (solvent B) was programmed as follows: starting at 10% B at 0.0 min, linearly increasing to 95% B by 15.0 min, holding at 95% B from 15.0 to 19.0 min, followed by re-equilibration at 10% B for 3.0 min. The flow rate was maintained at 0.3 mL/min, and the column temperature was set to 30 °C. The HPLC elute was introduced to the mass spectrometer through ESI.

### PCA analysis

Data analysis was performed in Python (v3.12.11) using Google Colab. Raw metabolomic data were imported from CSV format and processed using pandas and numpy. Preprocessing included removal of a header-like row, conversion of sample columns to numeric format, and exclusion of entries with missing identifiers. Sample names were standardized and indexed to ensure uniqueness.

Metabolite intensity values were normalized using z-score transformation with StandardScaler. Principal Component Analysis (PCA) was conducted using sklearn.decomposition.PCA, and the components were visualized with seaborn and matplotlib. Sample clustering was assessed across four experimental groups: Stationary_Horizontal, Stationary_Vertical, Rotational_Horizontal, and Rotational_Vertical. The proportion of variance explained by PC1 and PC2 was annotated.

Welch’s t-test (ttest_ind) was used to compare metabolite abundance between groups. For each feature, log_2_ fold change and –log_10_(p-value) were calculated. Volcano plots were generated using seaborn, with significance thresholds set at *p < 0.05, p < 0.01*, and *p < 0.001*. Features were considered significant if p-values were below the threshold and absolute log_2_ fold change exceeded 1. Significant and non-significant features were exported as separate CSV files.

Annotated plots included fold-change and p-value cutoffs, with labeled features meeting both criteria.

## Results

### Metabolites identification by LC-MS/MS

UPLC-MS/MS was utilized to identify and quantify the metabolites in the kale samples. First, the extraction solvent was tested by comparison of the intensities/area of the compound peaks generated. 70% Methanol in water extract gave larger areas for the majority of the compounds than the methanol and absolute ethanol extracts. Thus, 70% methanol is chosen for extraction.

The compounds were identified by the combination of searching of six libraries, including MassList, ChemSpider, Metabolika, mzVault, mzCloud, and Predicted Compositions.

### Impact of metabolites by microgravity

The PCA score plot (Figure 1) revealed distinct clustering among the four experimental conditions. SHS samples formed a compact group clearly separated from SVS along PC1, indicating that static horizontal orientation used to mimic microgravity produced a measurable shift in global metabolite profiles. RHS samples were positioned apart from RVS, showing partial but consistent separation along both PC1 and PC2. Both PC1 (38.6 %) and PC2 (27.6 %) accounted for 66.2 % of the total variance, demonstrating that orientation and rotational movement are major sources of metabolic variability in the dataset.

**Figure 1.**
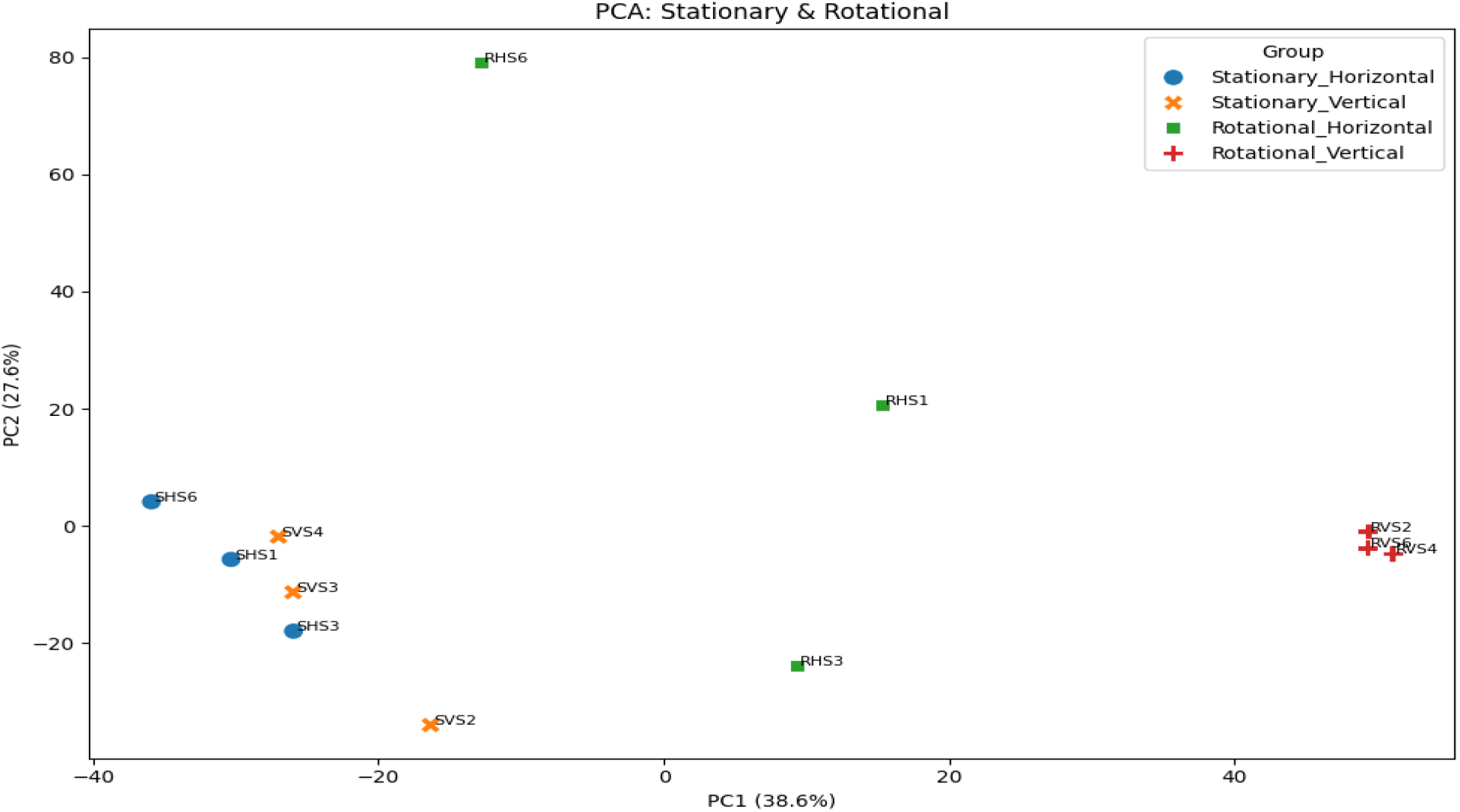
PCA score plot of metabolomic profiles under simulated microgravity. Samples were grouped by orientation and motion: Stationary Horizontal (SHS, blue circles), Stationary Vertical (SVS, orange crosses), Rotational Horizontal (RHS, green squares), and Rotational Vertical (RVS, red pluses). The first two principal components (PC1 and PC2) explain 38.6 % and 27.6 % of the total variance, respectively, providing a cumulative 66.2 % of explained variance.

Differential metabolite expression between RHS and RVS groups revealed a broad distribution of fold changes (Figure 2). Numerous metabolites (red dots) exceeded the *p < 0.05* threshold, and many of these also crossed the ±1 log_2_ fold-change cut-offs, indicating biologically meaningful differences in abundance. Several highly significant metabolites were annotated near the top of the plot, reflecting both strong statistical support and large magnitude of change. The spread of significant points on both sides of the vertical thresholds demonstrates that horizontal rotation under simulated microgravity induced both up-regulation and down-regulation of specific metabolites compared with vertical rotation.

**Figure 2.**
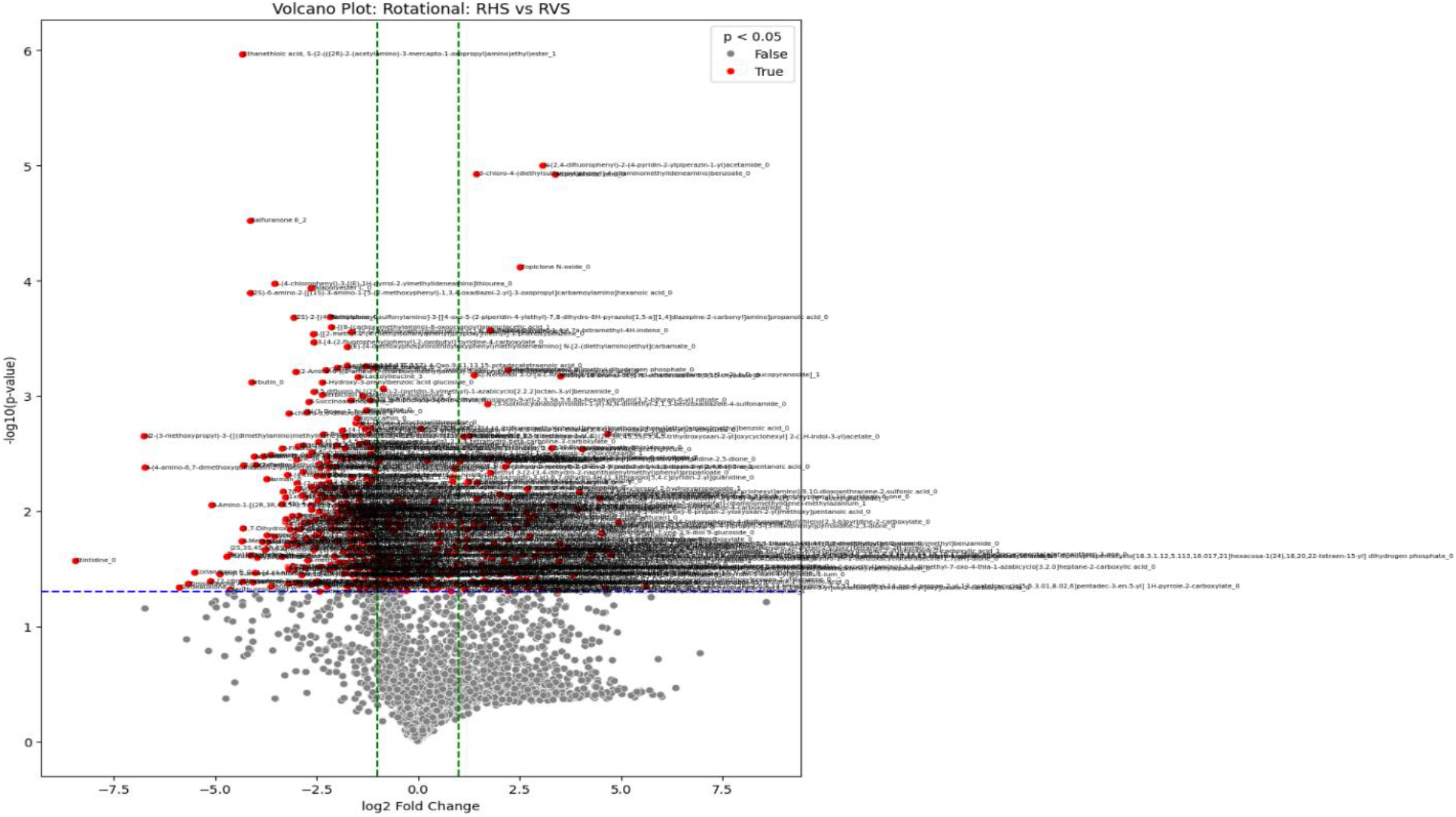
Volcano plot comparing metabolite profiles of RHS versus RVS samples under simulated microgravity. The x-axis shows log_2_ fold change in metabolite abundance; the y-axis displays – log_10_(p-value). Red points indicate metabolites with *p < 0.05* (significant), while grey points are not significant. Vertical green dashed lines represent ±1 log_2_ fold-change thresholds and the horizontal blue dashed line indicates the p = 0.05 significance cutoff.

The volcano plot in Figure 3 highlights a broad set of metabolites whose abundance differs between SHS and SVS samples. Numerous features (red points) surpass the *p < 0.05* cutoff, and many exceeded the ±1 log_2_ fold-change boundaries, indicating that horizontal orientation under stationary conditions induced pronounced metabolic shifts. Significant metabolites were distributed on both sides of the vertical thresholds, revealing both up-regulated and down-regulated compounds in the horizontal orientation relative to the vertical control.

**Figure 3.**
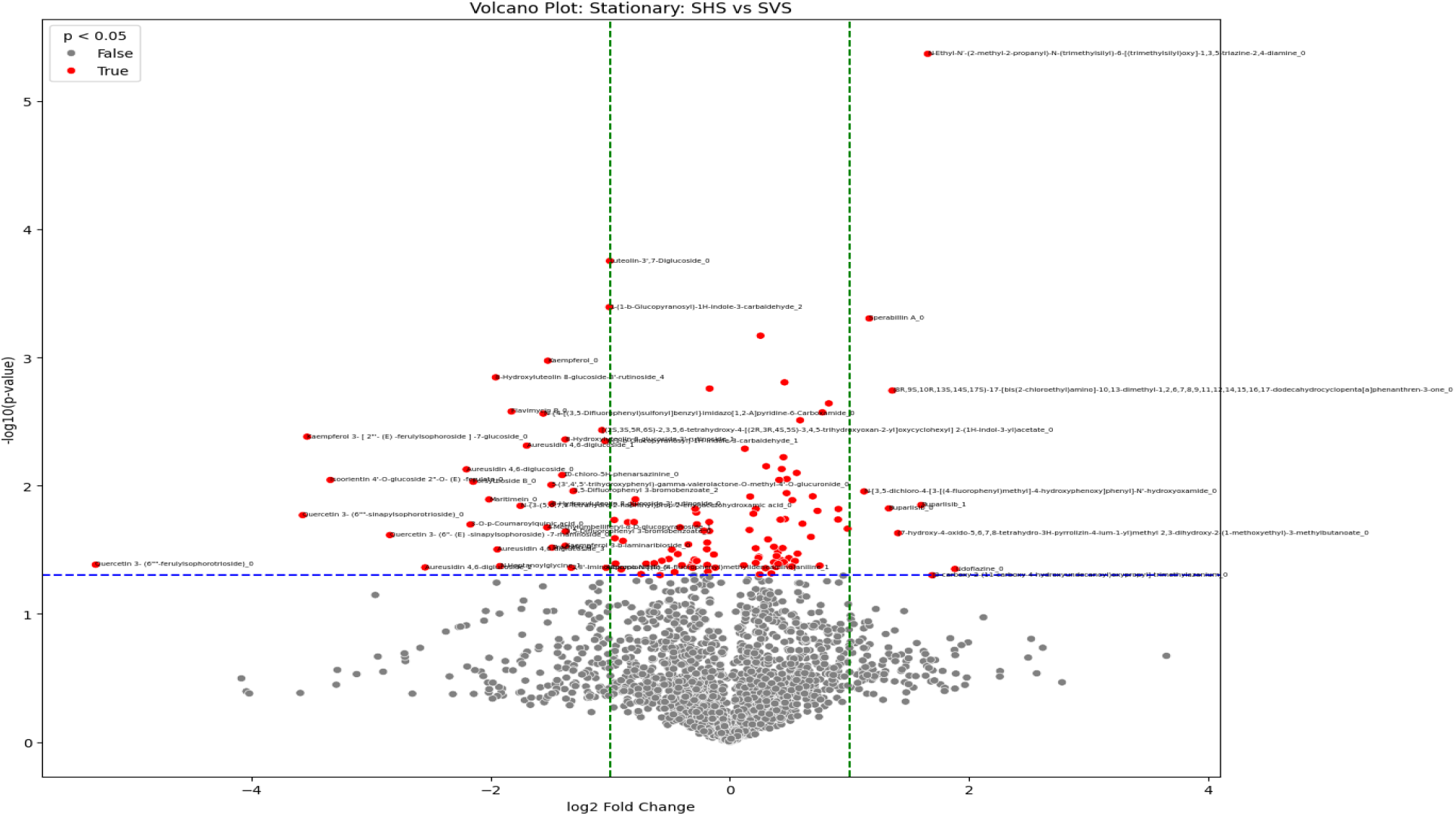
Volcano plot comparing metabolite profiles between Stationary Horizontal (SHS) and Stationary Vertical (SVS) samples. The x-axis shows the log_2_ fold change in metabolite abundance; the y-axis represents –log_10_(p-value). Red points indicate metabolites with *p < 0.05*, whereas grey points were not significant. The green dashed vertical lines mark the ±1 log_2_ fold-change limits, and the blue dashed horizontal line denotes the *p = 0.05* significance threshold

Applying a more stringent *p < 0.001* filter sharply reduced the number of significantly altered metabolites (Figure 4), isolating a high-confidence subset of features with large and statistically robust changes in abundance. Most of these metabolites exceeded the ±1 log_2_ fold-change limits, highlighting compounds that were both strongly and confidently modulated by horizontal orientation. The remaining non-significant features cluster around the center of the plot, reflecting background metabolic variation.

**Figure 4.**
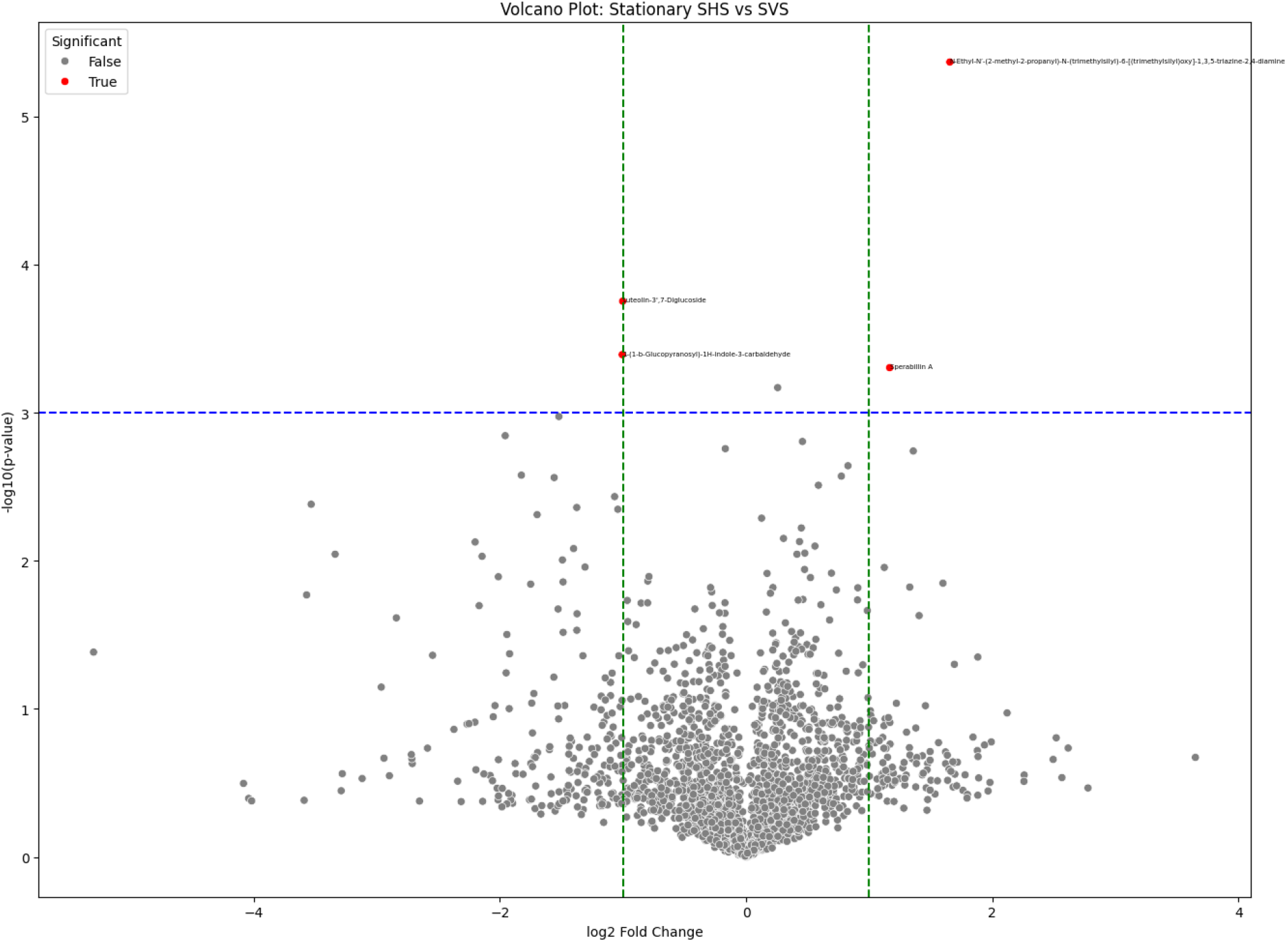
Volcano plot of metabolite differences between SHS and SVS samples at a stringent significance level (*p < 0.001*). The x-axis shows log_2_ fold change in metabolite abundance, and the y-axis shows –log_10_(p-value). Red points represent metabolites meeting the *p < 0.001* cutoff, while grey points were not significant. Green dashed vertical lines denote the ±1 log_2_ fold-change thresholds and the blue dashed horizontal line marks the *p = 0.001* boundary.

At the stricter *p < 0.001* threshold, only a few metabolites remained significantly different between RHS and RVS samples (Figure 5). These high-confidence features, highlighted in red, showed both large magnitude changes (beyond ±1 log_2_ fold change) and strong statistical support. Among them, key annotated metabolites including Ethanethoic acid, 5-[2-((2R)-2-(acetylamino)-3-mercapto-1-oxopropyl)amino]ethyl]ester and N-[2-(2,4-difluorophenyl)-2-(4-pyridin-2-ylpiperazin-1-yl)acetamide, displayed pronounced up-regulation in horizontal rotation and vertical rotation.

**Figure 5.**
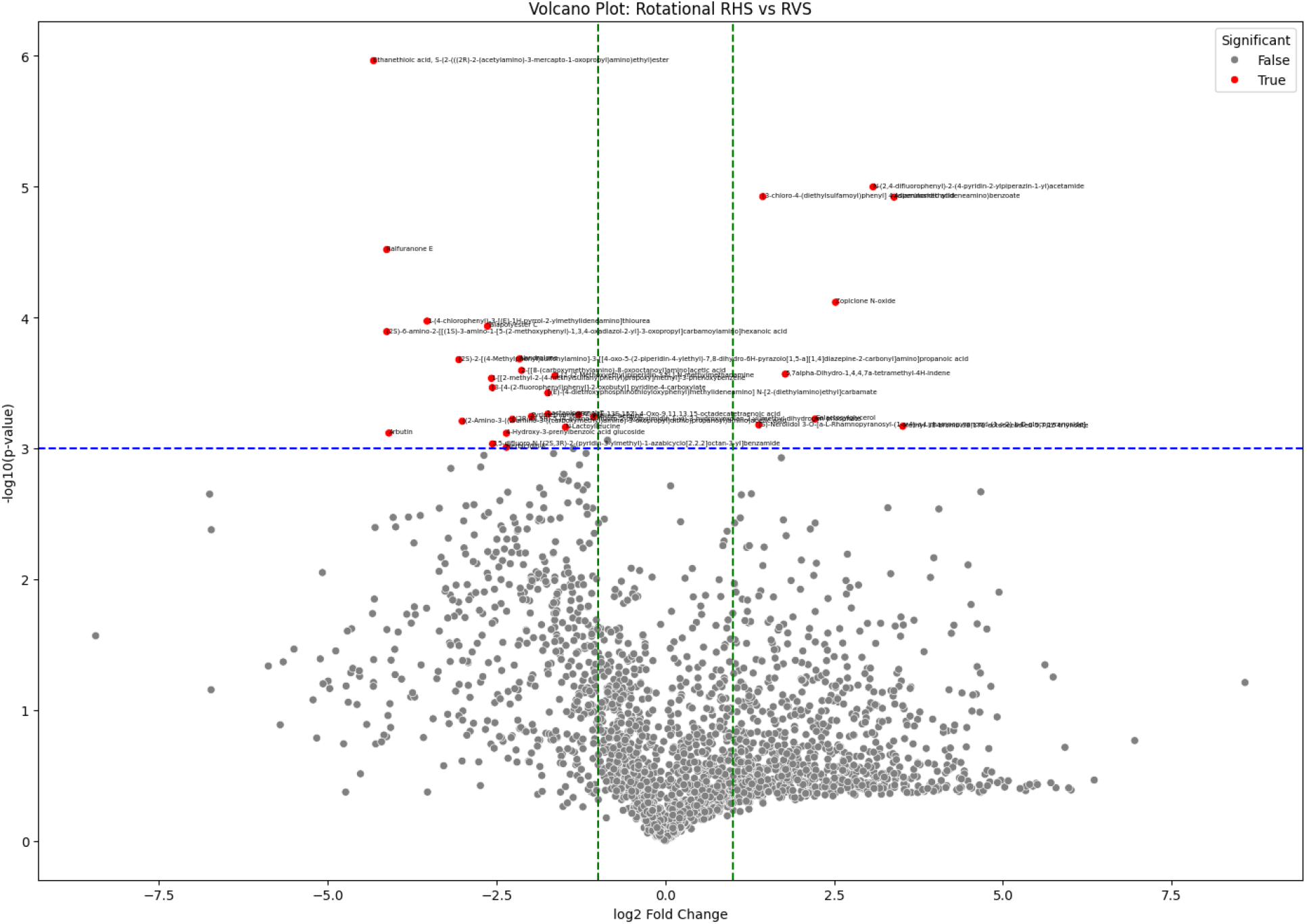
Volcano plot comparing metabolite abundance between Rotational Horizontal (RHS) and Rotational Vertical (RVS) samples at a stringent significance threshold (*p < 0.001*). The x-axis shows the log_2_ fold change, while the y-axis represents –log_10_(p-value). Red points mark metabolites meeting the *p < 0.001* cutoff; grey points wer not significant. Vertical green dashed lines indicate the ±1 log_2_ fold-change boundaries and the horizontal blue dashed line represents the *p = 0.001* significance level.

At the intermediate significance threshold of *p < 0.01*, the volcano plot (Figure 6) highlights a distinct subset of metabolites that were differentially expressed between SHS and SVS samples. Several red points exceed both the p-value cut-off and the ±1 log_2_ fold-change limits, indicating metabolites with both substantial and statistically reliable changes in abundance. These features represent a refined, yet broader set compared with the *p < 0.001* analysis, capturing both robust and moderately significant alterations caused by horizontal orientation under stationary conditions.

**Figure 6.**
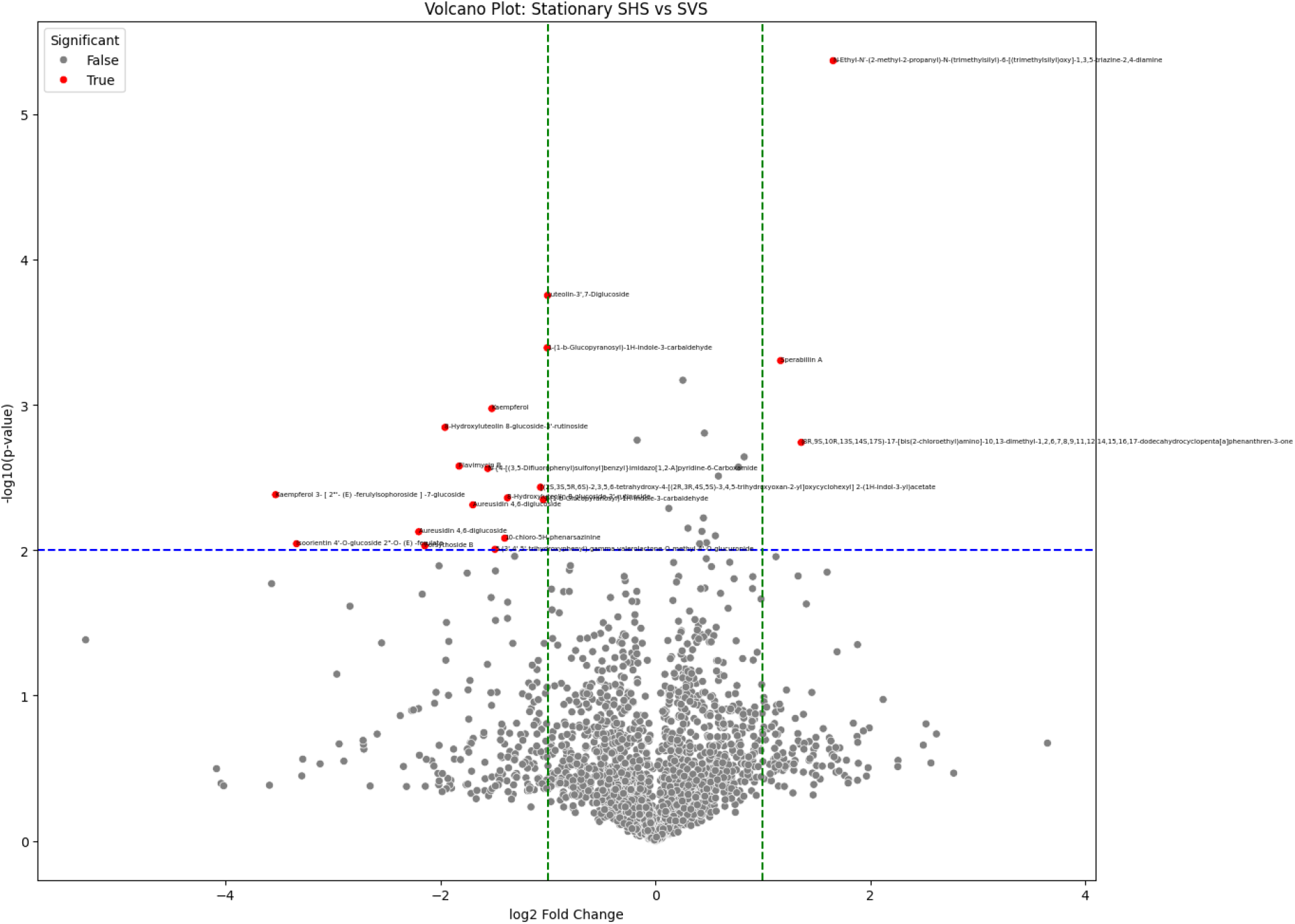
Volcano plot of metabolite abundance differences between SHS and SVS samples at *p < 0.01*. The x-axis displays log_2_ fold change, while the y-axis shows –log_10_(p-value). Red points represent metabolites significant at *p < 0.01*; grey points were not significant. Vertical green dashed lines mark the ±1 log_2_ fold-change thresholds and the horizontal blue dashed line indicates the *p = 0.01* significance level.

At the intermediate *p < 0.01* threshold, the volcano plot (Figure 7) highlights a defined set of metabolites significantly altered by horizontal rotation compared with vertical rotation. Several red points surpass both the statistical and fold-change criteria, indicating robust differences in metabolite abundance. Notably, key annotated metabolites including Ethanethoic acid, 5-[2-((2R)-2-(acetylamino)-3-mercapto-1-oxopropyl)amino]ethyl]ester showed marked up-regulation in horizontal rotation, suggesting that rotational microgravity produces pronounced metabolic modulation beyond that of vertical controls.

**Figure 7.**
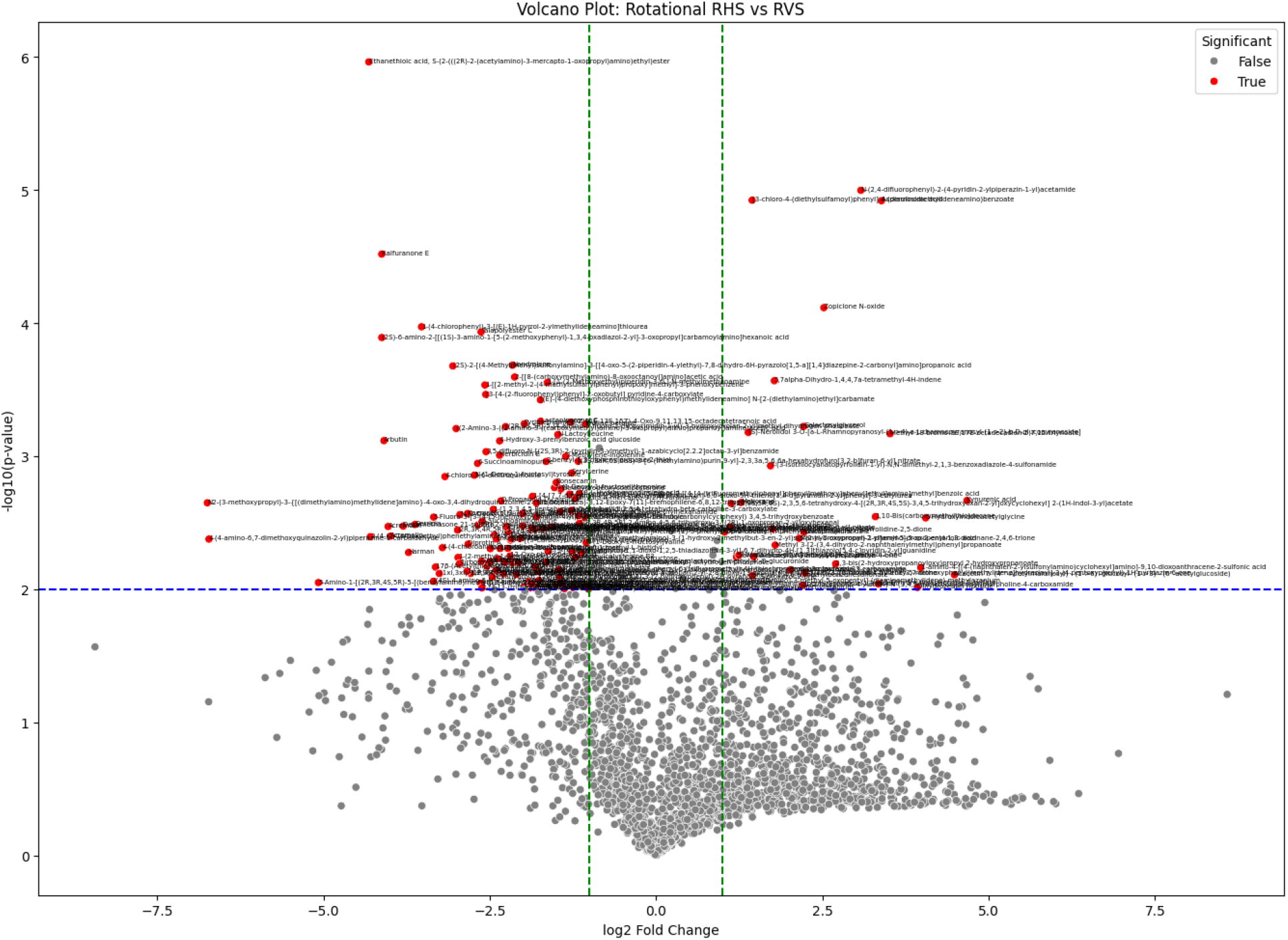
Volcano plot showing differentially expressed metabolites between RHS and RVS samples at a *p < 0.01* significance threshold. The x-axis indicates log_2_ fold change, while the y-axis represents –log_10_(p-value). Red points denote metabolites significant at *p < 0.01*; grey points are not significant. The vertical green dashed lines mark ±1 log_2_ fold-change boundaries, and the horizontal blue dashed line indicates the *p = 0.01* significance level.

## Discussion

The clear separation between SHS and SVS clusters supports the hypothesis that a horizontal, microgravity-like orientation disturbed cellular metabolism even in the absence of motion. This observation aligned with previous reports that microgravity simulation induces widespread metabolic reprogramming, including changes in energy metabolism and stress-response pathways (Herranz et al., 2013; Horneck et al., 2010). The partial segregation of RHS and RVS groups suggested that dynamic microgravity simulation (rotation) imposes additional metabolic pressures, which is consistent with the findings of Nickerson et al. (2000), that mechanical forces and fluid shear in rotating bioreactors further influence metabolite fluxes. The two principal components captured over two-thirds of the total variance, indicating that orientation and rotational cues were dominant drivers of metabolic divergence in this system. Such distinct clustering supports the sensitivity of cellular metabolic networks to gravitational and mechanical stimuli and highlights PCA as an effective tool for visualizing these global shifts (Demšar et al., 2013).

The volcano plot highlights a robust set of metabolites whose abundance is significantly altered by horizontal rotational orientation, reinforcing the PCA observation that rotational microgravity imposes distinct metabolic pressures (Zhou et al., 2025). Metabolites with large positive or negative fold changes likely represent key nodes in pathways sensitive to gravitational or mechanical cues. Similar orientation-dependent metabolic changes have been reported in previous microgravity studies, where altered fluid shear and gravitational forces modulate cellular energy metabolism, oxidative stress responses, and signaling networks (Wei et al., 2022; St-Martin et al., 2024; Chen et al., 2025). The presence of both up- and down-regulated metabolites suggested a complex regulatory response, possibly involving compensatory adjustments in primary metabolic pathways such as amino acid and lipid metabolism. Identifying these metabolites and mapping them to biochemical pathways will be important for elucidating mechanisms of microgravity adaptation and for designing countermeasures for spaceflight-related metabolic disturbances.

These findings demonstrate that static horizontal positioning used as a microgravity analogue elicits robust metabolic reprogramming even in the absence of rotational movement. Such orientation-dependent changes in metabolite abundance were consistent with recent reports that microgravity alters core biochemical processes, including energy metabolism and redox balance (Han et al., 2024; St-Martin et al., 2024; Serrano & Medina, 2025). The presence of both increased and decreased metabolites suggested a complex adaptive response, likely involving multiple metabolic pathways.

The focused set of metabolites identified at *p* < 0.001 provided stronger evidence that static horizontal positioning used to simulate microgravity induced profound metabolic reprogramming. Similar reductions in significant features under more conservative thresholds have been observed in recent microgravity metabolomics studies, indicating a core group of gravity-sensitive metabolites that remain significant even after stringent correction (Thiel et al., 2021; Ibrahim et al., 2024). These robust changes likely represent primary metabolic pathways most responsive to altered gravitational cues, such as energy metabolism, oxidative stress responses, and signaling cascades.

This reduced but well-defined set of metabolites underscores the robustness of the metabolic response to dynamic microgravity simulation. Even after stringent multiple-testing control, horizontal rotation continued to induce significant shifts, reinforcing the notion that mechanical cues and rotational microgravity exert distinct biochemical effects. Similar high-confidence metabolite signatures have been reported in recent omics studies of microgravity, where shear stress and altered gravitational forces drive persistent changes in energy metabolism and stress-response pathways (Sharma & Curtis, 2022; Lozzi et al., 2025). The identification of a small cluster of highly significant metabolites provides a focused starting point for downstream pathway analysis and the discovery of potential biomarkers of gravitational adaptation.

The metabolites identified at *p* < 0.01 provided a balance view of orientation-induced metabolic changes, capturing both high-confidence and moderately significant shifts. This intermediate filter reinforces evidence that static horizontal orientation used as a microgravity analogue drives significant reprogramming of metabolic networks. Similar patterns of orientation-dependent metabolic modulation have been observed in recent omics studies of real and simulated microgravity, where energy metabolism, oxidative stress responses, and signaling pathways were commonly affected (Nguyen et al., 2021; Abdelfattah et al., 2024). The findings suggest that horizontal orientation alone, even without rotational motion, is sufficient to trigger biochemical responses characteristic of microgravity adaptation.

These findings reinforce the evidence that rotational microgravity exerts measurable biochemical effects on cellular metabolism. The intermediate *p* < 0.01 threshold captures both strongly and moderately significant changes, offering a balanced view of metabolic adaptation compared with the more stringent *p* < 0.001 analysis. Similar orientation-dependent metabolic responses have been documented in recent omics investigations of real and simulated microgravity, where altered fluid shear and gravitational forces drive shifts in energy metabolism and stress-related pathways (Thiel et al., 2021; St-Martin et al., 2024; Bergstrom et al., 2025). The identification of these significantly altered metabolites provides a valuable foundation for pathway analysis and for developing potential biomarkers of microgravity-induced metabolic reprogramming.

## Conclusion

Simulated microgravity, achieved through both static horizontal positioning and horizontal rotation, induced clear and statistically robust shifts in metabolite profiles, highlighting gravity-responsive metabolic pathways and potential biomarkers that can guide future studies and the development of spaceflight countermeasures.

## Acknowledgements

The authors acknowledge support from NASA (Grant # 80NSSC22K0869).

